# The sudden emergence of a *Neisseria gonorrhoeae* strain with reduced susceptibility to extended-spectrum cephalosporins, Norway

**DOI:** 10.1101/2020.02.07.935825

**Authors:** Magnus N. Osnes, Xavier Didelot, Jolinda de Korne-Elenbaas, Kristian Alfsnes, Ola B Brynildsrud, Gaute Syversen, Øivind Nilsen, Birgitte F. de Blasio, Dominique A Caugant, Vegard Eldholm

**Affiliations:** Division of Infection Control and Environmental Health, Norwegian Institute of Public Health, Oslo, Norway; School of Life Sciences and Department of Statistics, University of Warwick, Coventry, United Kingdom; Department of Infectious Diseases, Public Health Service Amsterdam, Amsterdam, The Netherlands; Department of Microbiology, Oslo University Hospital Ullevaal, Oslo, Norway; Oslo Centre for Biostatistics and Epidemiology, Department of Biostatistics, Institute of Basic Medical Sciences, University of Oslo, Oslo, Norway; Department of Community Medicine, Faculty of Medicine, University of Oslo, Oslo, Norway; The AMR Centre, Norwegian Institute of Public Health, Oslo, Norway

## Abstract

The *Neisseria gonorrhoeae* multilocus sequence type (ST) 7827 emerged in dramatic fashion in Norway in the period 2016-2018. Here, we aim to determine what enabled it to establish and spread so quickly. In Norway, ST-7827 isolates were almost exclusively isolated from men. Phylogeographic analyses demonstrated an Asian origin of the ST with multiple importation events to Europe. The ST was uniformly resistant to fluoroquinolones and associated with reduced susceptibility to both azithromycin and the extended-spectrum cephalosporins (ESC) cefixime and ceftriaxone. We identified additional independent events of acquisition of *penA* and *porB* alleles in Europe, associated with further reduction in cefixime and ceftriaxone susceptibility, respectively. Transmission of the ST was largely curbed in Norway in 2019, but our results indicate the existence of a reservoir in Europe. The worrisome drug resistance profile and rapid emergence of ST-7827 calls for close monitoring of the situation.

## Introduction

*Neisseria gonorrhoeae* represents an increasing burden on health (1). Globally, the pathogen has acquired resistance to all drugs recommended for therapy (2). As a result of the rapid acquisition of antibiotic resistance (ABR) and accompanying increased risk of treatment failure, the World Health Organization recently listed *N. gonorrhoeae* as a high-priority pathogen, on which research and drug development should focus (3).

The incidence of gonorrhea is increasing rapidly in Norway. From 1995 to 2010, the number of cases fluctuated between 150-300 cases per year, but has since followed a steep upward trajectory, reaching 1704 reported cases in 2019 (4). Of these cases, 78% were men and 58% were infected following homosexual sex (4).

Gonococci are efficient colonizers of mucosae, and can in addition to the urethrae, infect both the pharynx and rectum. In men, urethral infections are mostly symptomatic, and affected individuals tend to seek treatment rapidly, whereas pharyngeal and rectal infections often are asymptomatic and may go undetected for a longer period (5). In women, cervicitis is the most common manifestation, and infections are often asymptomatic. It is thus possible that different infection sites play differential roles in the epidemic process.

Genome-based analyses of geographical dispersal and person-to-person transmission have the potential to uncover unobserved drivers of an epidemic. Previously, such methods have been used to determine time periods for which an outbreak has been poorly sampled (6), declare the end of an outbreak (7), estimate the geographical origin of extant gonorrhea (8) and assess the importance of importation for outbreak sustenance (9). However, obtaining representative samples is often a challenge when reconstructing phylogeographic and transmission histories.

In the period 2016-2018, there was a sudden surge in cases belonging to the hitherto undescribed sequence type (ST) 7827 among men in Norway. The number of cultured cases belonging to the ST rose from 11 (2.9%) in 2016 to 124 (16.6%) of cultured cases in 2018. Here, we apply genome-based analyses to investigate the sudden emergence of ST-7827.

## Materials and methods

### Clinical isolates

The Norwegian Institute of Public Health hosts the National Reference Laboratory for Gonorrhea and receives all cultured cases from Norway. Whole-genome sequencing has been performed routinely since 2016. Multi-locus sequence typing (MLST) is performed *in silico* (10) against the pubMLSTdatabase (11) to determine STs. In the period 2016-2018 159 isolates were found to belong to ST-7827. Illumina short-read data were retrievable for 148. Out of 147 isolates with associated gender information, 143 were sampled from men. Sequence data are available under study accession PRJEB32435. Beyond the study period, MLST results were retrieved from our strain collection for the year 2019.

In addition, 24 ST-7827 isolates with available sequence data were identified in the PubMLST database (11) and four assemblies were identified and downloaded from PathogenWatch (12). Finally, we screened the metadata attached to available large genome datasets uploaded to the NCBI short read archive (https://www.ncbi.nlm.nih.gov/sra) in order to identify additional ST-7827 isolates. For datasets where this information was lacking, we downloaded and screened the sequence data *in silico* against the pubMLST database to determine the ST. We did not identify any ST-7827 members among recently published sequences from Japan (13), the USA (14) or a global gonorrhea genomic study (8). However, in a recent study from China, 74 out of 435 isolates belonged to ST-7827 (15), and these were included in the current study. Information on all isolates is available in a supplementary excel file (Appendix 2: supplementary_<u>isolate_data</u>). The current project was approved by the Norwegian Regional Ethical Committee (project identifier 2019/347).

### Genome analyses

Raw Illumina reads were assembled *de novo* using SPAdes (16) in «careful» mode. The assemblies were further filtered to remove contigs with a kmer-coverage < 3 and length < 500 nucleotides.

To generate a high-quality reference genome for comparative genomic analyses, long read data were generated on the Oxford Nanopore GridION platform for one of the Norwegian isolates (641189). DNA was extracted and sequences generated as described previously (17). For assembly we employed a hybrid approach implemented in Unicycler v0.4.7 (18), leveraging both long Oxford Nanopore reads and short accurate Illumina reads. This resulted in a resolved circular genome of 2,215,865 bp plus a 4,207 bp plasmid.

Parsnp (19) was used to align the short read assemblies to the closed reference genome (excluding the plasmid). A whole-genome multifasta retaining the reference nucleotide position for all sites by filling all non-core regions with reference nucleotides was generated with an in-house script (https://github.com/krepsen/parsnp2fasta).

Internal repeat regions in the reference genome were identified using Mimeo (20). A minimum length of 200 nucleotides with ≥ 95% identity was required for repetitive regions to be called. A total of 134,107 sites were identified in the reference and masked from all sequences contained in the whole-genome multifasta using the EMBOSS (21) packages seqretsplit and maskseq.

Finally, Gubbins (22) was employed to identify recombination tracts in the whole-genome alignments and a recombination-corrected phylogeny was generated as per default by Gubbins. To root the phylogeny we performed a separate analysis including an ST-1901 genome.

### Analyses of antibiotic susceptibility

Minimum inhibitory concentration (MIC) data were available for a handful of the isolates downloaded from PubMLST and PathogenWatch. For the Norwegian isolates, full MIC profiles were available for all isolates from 2016-2017, and about 80% of 2018 isolates. Throughout the text, we refer to EUCAST (23) when breakpoints are discussed (Resistance breakpoints: cefixime and ceftriaxone > 0.125 µg/ml; ciprofloxacin > 0.06 µg/ml). Until recently, azithromycin MIC ≥ 0.5 µg/ml were considered as intermediately resistant by EUCAST, whereas the latest guideline refrains from defining breakpoints. Here we use the term “reduced susceptibility” on isolates with azithromycin MICs of ≥ 0.25 µg/ml and cefixime/ceftriaxone MICs of ≥ 0.064 µg/ml.

Mutations and alleles of ABR genes other than *penA* were identified in PathogenWatch with the AMR <u>Library 485 version 0.0.1</u> (24). To annotate *penA* alleles, genome assemblies were aligned against the *penA* allele database downloaded from PubMLST (11), using BLAST (25). When available, the NGSTAR (26) allele nomenclature was used. Results from phenotypic and *in silico* predicted resistance are available as supplementary data (Appendix 2: supplementary_<u>isolate_data</u>). Identified resistance determinants from PathogenWatch are not included in Fig. 1 but can be found annotated on the phylogeny in Fig. S4.

**Figure 1.**
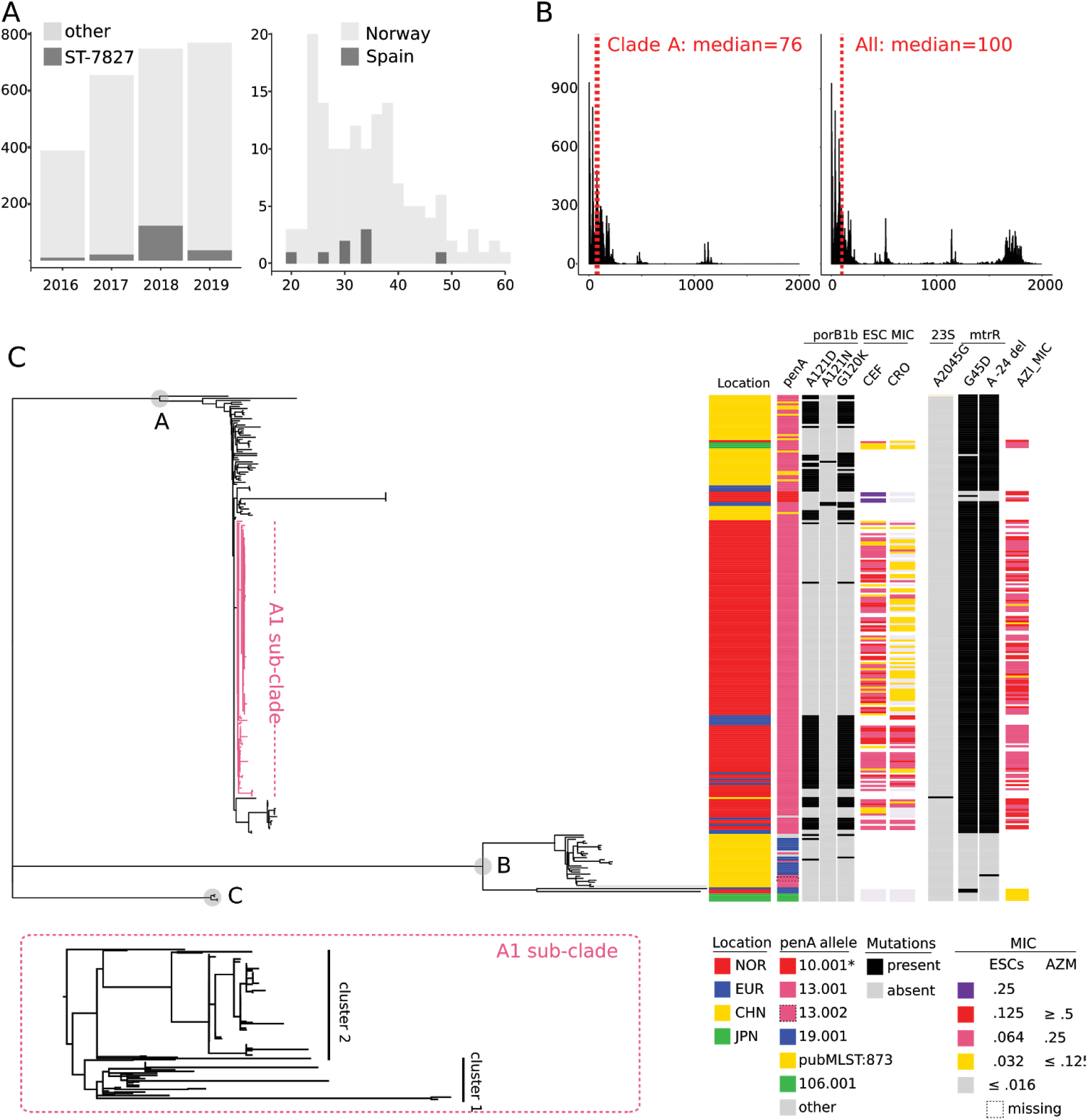
Overview of ST-7827 incidence and genomic properties. A: Incidence of ST-7827 among culture-positive *N. gonorrhoeae* cases in relation to the total number of cultured cases 2016-2019 (left). Age distribution of ST-7827 cases in Norway and Spain (right) B: Pairwise SNP-distances within Clade A and the entire ST-7827 collection respectively. C: Annotated recombination-corrected genome-based ST-7827 phylogeny. NOR=Norway, EUR=Europe, CHN=China, JPN=Japan, ESCs = extended spectrum cephalosporins (cefixime, ceftriaxone), AZM = azithromycin. Due to the low number of samples, isolates from the United Kingdom, Spain, Italy and France were combined in a single “Europe” category. The European A1 sub-clade is highlighted in dark pink. A more detailed view of the A1 phylogeny is shown on the bottom.

### Additional phylogenetic analyses and transmission modeling

Phylogeographic inferences were made using stochastic character mapping (27) implemented in the phytools R package (28). To account for phylogenetic uncertainty we constructed a set of 100 bootstrap trees with PhyML (29) using the polymorphic sites estimated to be non-recombinant according to Gubbins. For each of the bootstrap trees, we simulated 10 stochastic character mappings, giving us a posterior set of 1000 geographically mapped phylogenetic trees, from which a consensus tree was generated.

The Gubbins output was used directly as input for BactDating (30) to perform root-to-tip regression and temporal analyses. TransPhylo (6) was used to estimate transmission trees. For the generation time distribution (time from being infected to infecting others), we considered two gamma distributions, one which was estimated in an individual-based modeling study from a community of men who have sex with men (MSM) (31) - “prior 1” with shape 0.57 and scale 0.30, and a distribution mimicking prior 1 but penalizing transmissions in the incubation phase - “prior 2” with shape 1.20 and scale 0.14 (Fig. S5). The sampling distribution (time between infection and sampling) was set equal to the generation time distribution. Based on the discrepancy between the number of reported cases and cultured cases over time, we know that about 55% of cases are lost due to failed culturing. In addition, we must assume that some, particularly asymptomatic cases, go undiagnosed. Yet, In Norway, this fraction is expected to be modest as high-risk individuals are screened frequently and contact tracing is performed on all cases. Based on the above, we fixed sampling densities in TransPhylo to the fractions 0.2, 0.3, 0.4 and 0.5 of the number of cases. Sensitivity testing indicated that the modeling output was not overly sensitive to the choice of generation time and sampling density priors (Fig. S8). In the end, we used the results generated with a sampling density of 0.4 and prior 2 for the generation time.

From the estimated transmission trees we extracted the posterior generation times, the number of secondary infections caused by primary infections, the pairwise individual-to-individual probability of direct transmissions and the average number of intermediate carriers between each pair of patients. To investigate whether infections of the urethra, rectum and pharynx differed in their propensity to transmit to secondary cases, we stratified the posterior generation times and the number of secondary infections, on different infection sites.

The methods applied in this section are described in detail in Appendix 1.

## Results

### Population-genomic characteristics of ST-7827

A total of 251 genomes belonging to ST-7827 were identified and included in the study, of which 148 were samples from Norway isolated in the study period January 2016 to December 2018, whereas the rest were from China, Japan and Europe (see Materials and methods). Among the Norwegian cases, 97% were men and 90% resided in the greater Oslo area. The proportion of ST-7827 increased abruptly from 2.9 % to 16.6 % of all isolates in 2016 and 2018 respectively, with most cases being adult men 25-50 years old (Fig. 1 A). Demographic data were also available for Spanish isolates. These were all sampled from men, whose age-distribution mirrored the one in Norway, indicating transmission in similarly structured MSM networks. Interestingly, in 2019, the transmission of the ST seems to have been largely interrupted in Norway, accounting for only 37 (4.8%) cultured cases.

The genome-wide SNP phylogeny revealed that ST-7827 consisted of three deep-branching clades (Fig. 1 B and C). We termed the largest clade containing the majority of isolates “Clade A” and a medium-sized clade “Clade B”. Four isolates from Japan belonged to a small separate clade termed “Clade C”. In the plot of pairwise distances, two dominating peaks representing inter and intra clade A and clade B distances can be seen together with two smaller peaks representing mainly Clade C versus clade A and B distances.

Most isolates belonging to Clade A were part of a tight sub-clade of European isolates bearing the hallmarks of recent spread (geographically restricted and short branch lengths), which we termed the A1 sub-clade. The majority of isolates in the A1 sub-clade were from Norway, but it also contained six Spanish and two Italian isolates.

All ST-7827 isolates except one harbored the *gyrA* S91F mutation and the vast majority of isolates carried additional *gyrA* and *parC* mutations. At the MIC level, all tested strains were resistant to ciprofloxacin (Appendix 2: supplementary<u>_isolate_data</u>). In Clade A, the *mtrR* promoter deletion −24A and genic G45D mutations were ancestral and nearly all tested isolates had azithromycin MICs between 0.25 and 1 µg/ml (Fig. 1 C).

The *penA* gene exhibited significant allelic diversity, with the 13.001 allele being dominant in Clade A. The *penA* 13.001 allele is non-mosaic but carries an A501V mutation and has been shown to be associated with moderately elevated ESC MICs of ≥ 0.03 µg/ml (32,33). Across the ST, primarily in Clade A, we also identified repeated acquisition of a *porB* allele carrying G120K and A121D mutations. These mutations have previously been indicated in elevated ESC MICs in combination with *penA (32)*. As can be seen in Fig. 1 C, acquisition of the mutated *porB* was associated with a reduction in ceftriaxone susceptibility, with the strains carrying it typically having MICs of 0.064 - 0125 µg/ml. The annotated phylogeny suggests that acquisitions of this *porB* allele have occurred both in Asia and Europe. A small cluster of cefixime resistant isolates had acquired a mosaic *penA* closely related to the 10.001, most likely in Europe (Fig. 1 C). The 10.001 allele and its variants have previously been shown to be associated with cefixime resistance, but not ceftriaxone (13), which fits our observations well. The allele identified here harbored two mutations (P538S and T549A) relative to 10.001, but was otherwise identical.

### Spatial and temporal inferences

From the phylogeny it is clear that the ancestor of ST-7827 emerged in Asia. To generate a more detailed understanding of the geographic origins and spread of Clade A, which was found to have expanded in Europe, we performed phylogeographic analyses employing SIMMAP (27) on this subset of isolates. A consensus tree of the estimated geographical states of the ancestral tree is shown in Fig. 2 A, and the estimated number of importation events from Asia to Europe (and vice versa) is shown in Fig. 2 B. Together, these figures indicate that Clade A was imported from East Asia (Japan, China or some unsampled east-Asian country) to Europe on multiple separate occasions. The estimated number of transition events between Norway and other geographical locations are shown in Fig. 2 C. Multiple introductions to Norway were inferred, with the most frequent transitions being between Norway and Spain.

**Figure 2.**
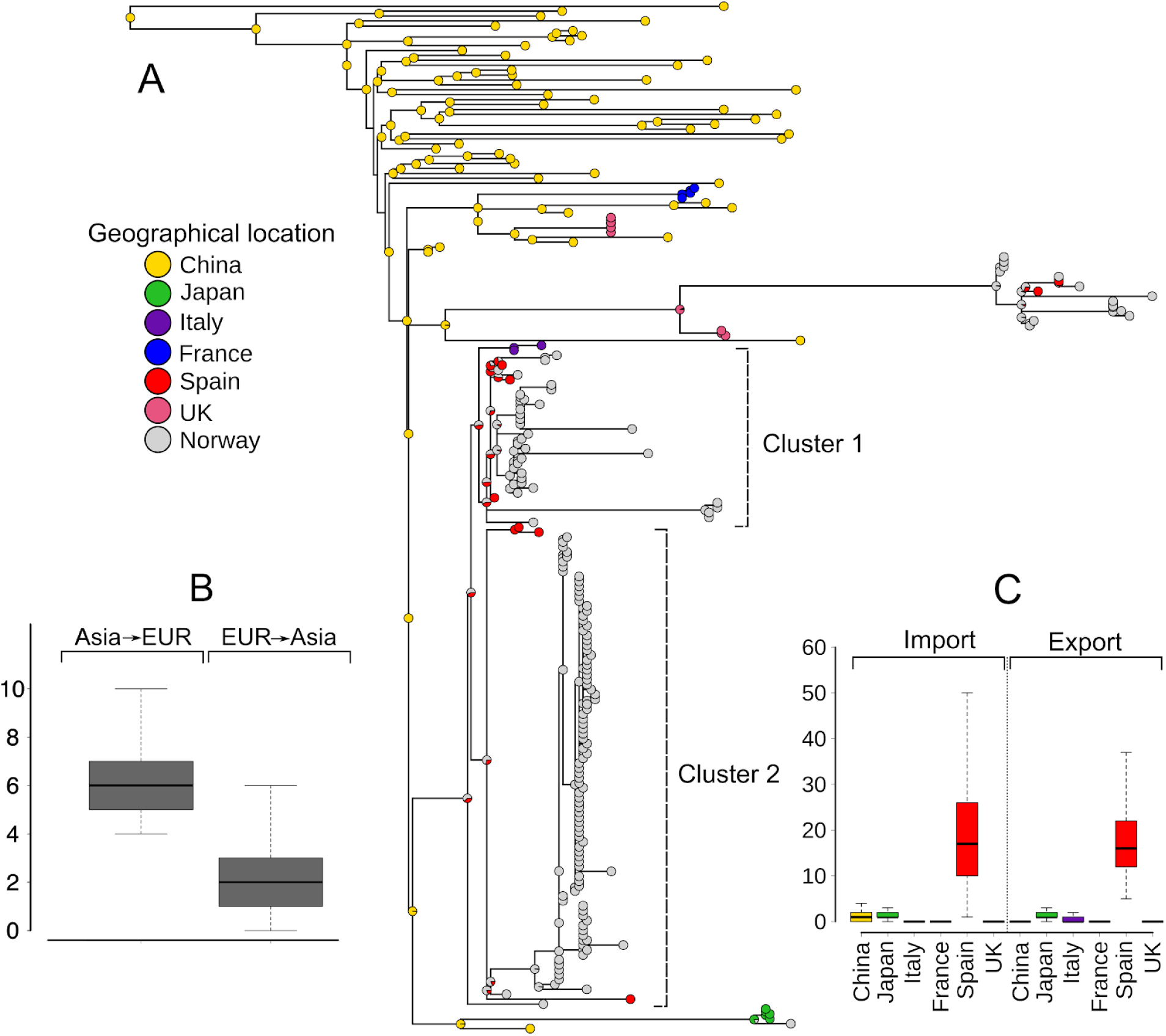
Phylogeographic inference. A: A consensus tree of the estimated geographical location at each node. Seven of the basal tips in Clade A (four from Japan and three from China) were removed from this plot because they had a very large genetic distance from the other samples which would collapse the branches and make the figure difficult to interpret. The internal nodes on all the removed branches were mapped to Asia. B: Box plot summarizing the realized number of transition events between Europe and Asia across 1000 SIMMAP simulations. C: Similar to B but for transition events from and to Norway.

A time-resolved phylogeny of the entire Clade A could not be generated due to a lack of temporal signal (see Supplementary Methods in Appendix 1). This was, however, possible for the much less diverse subset constituting the European sub-clade A1 (Fig. 3 A). The mean estimated substitution rate in the non-recombining regions of the genome was 2,5 × 10^−6^ per site per year (CI; 1,72 × 10^−6^ to 3,56 × 10^−6^), which is very similar to previous estimates *N. gonorrhoeae* mutation rates (8,14,34). Sub-clade A1 includes the majority of the sampled ST-7827 isolates (135/251) and contained two sub-clusters (Clusters 1 and 2) which diverged around 2008 (CI; 1997,5 to 2012,5) (Fig. 3 A). The phylogeographic analyses (Fig. 2 A) revealed that these clusters were introduced to Europe from Asia, as the geographical location is mapped to Asia in their common basal branches. The samples from Spain and Italy are placed basally within each cluster, suggesting that multiple introductions to Norway were followed by local transmission.

**Figure 3.**
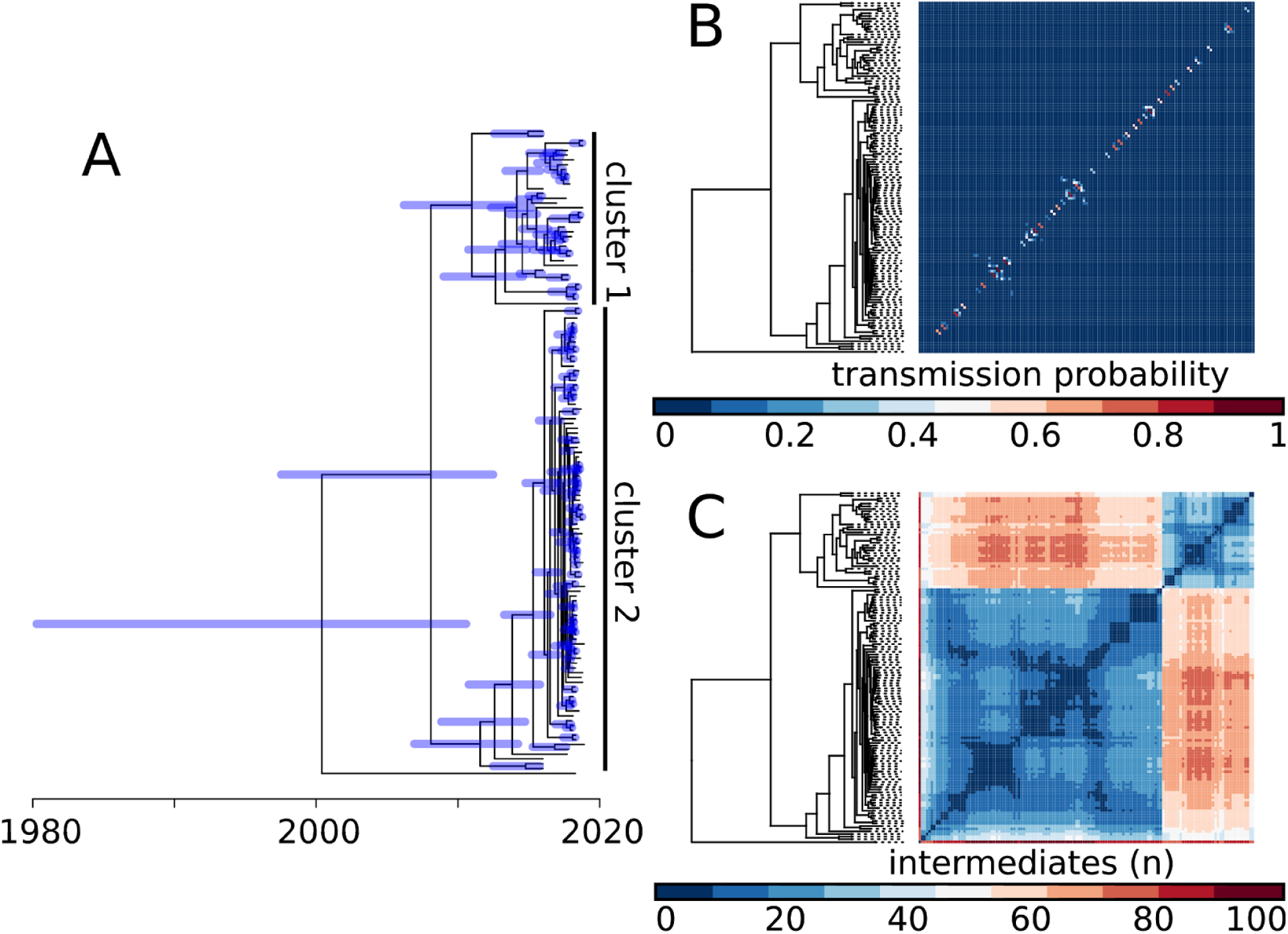
Reconstruction of temporal evolution and transmission patterns in the European outbreak. A: Time-stamped phylogeny of the European outbreak generated with BactDating. Node bars indicate 95% highest posterior density intervals. B: Heatmap ordered after the phylogeny showing the posterior probability of having a direct transmission between each pair of patients, calculated as the fraction of the transmissions trees where the pair is directly connected. C: Similar heatmap to B, showing the mean number of intermediate individuals between each pair of patients over all the transmission trees.

### Transmission analyses

In order to investigate transmission dynamics in more detail, we reconstructed genome-based transmission trees employing TransPhylo (6), with the time-stamped phylogeny as input. For the outbreak as a whole, the reproductive number was estimated to be 1.20 (credibility interval: 1.10, 1.31), and this was not sensitive to different assumptions on sampling densities (see Fig. S7 and Appendix 1). The estimated probabilities of direct transmission between each pair of patients are shown in Fig. 3 B. In total, there were 24 pairs with a direct transmission probability higher than 50% (cumulatively 33 with a probability higher than 40% and 42 higher than 30%). This shows that using only genomic data from the samples that are successfully grown Norway, we are not able to fully resolve the transmission chains, since for most patients there is a higher probability that their infector is unobserved than among the samples. The mean number of intermediate carriers is shown in Fig. 3 C.

Although intermediate unsampled carriers were inferred between most pairs of patients, groups of patients with few intermediates between them are clearly visible (note the blue squares in the heatmap which indicates very few intermediate carriers for the group as a whole). A consensus transmission tree is shown in Fig. 4 A. The smaller Cluster 1 contains samples from both Spain and Italy internally, while the larger Cluster 2 contains only Spanish samples on one of the basal branches. This suggests that Cluster 1 has been sustained by multiple importation events. For Cluster 2, the analyses point to a single introduction event from southern Europe, which then established itself in Norway and has been sustained by local transmission. Also of note is the observation that the strains belonging to Cluster 1, which have been imported repeatedly to Norway from Europe, carry a *porB* allele (G120K and A121D) associated with elevated ceftriaxone MICs (Fig. S3). This finding is consistent with ongoing circulation of ST-7827 in Europe with a worrisome drug-susceptibility profile.

**Figure 4.**
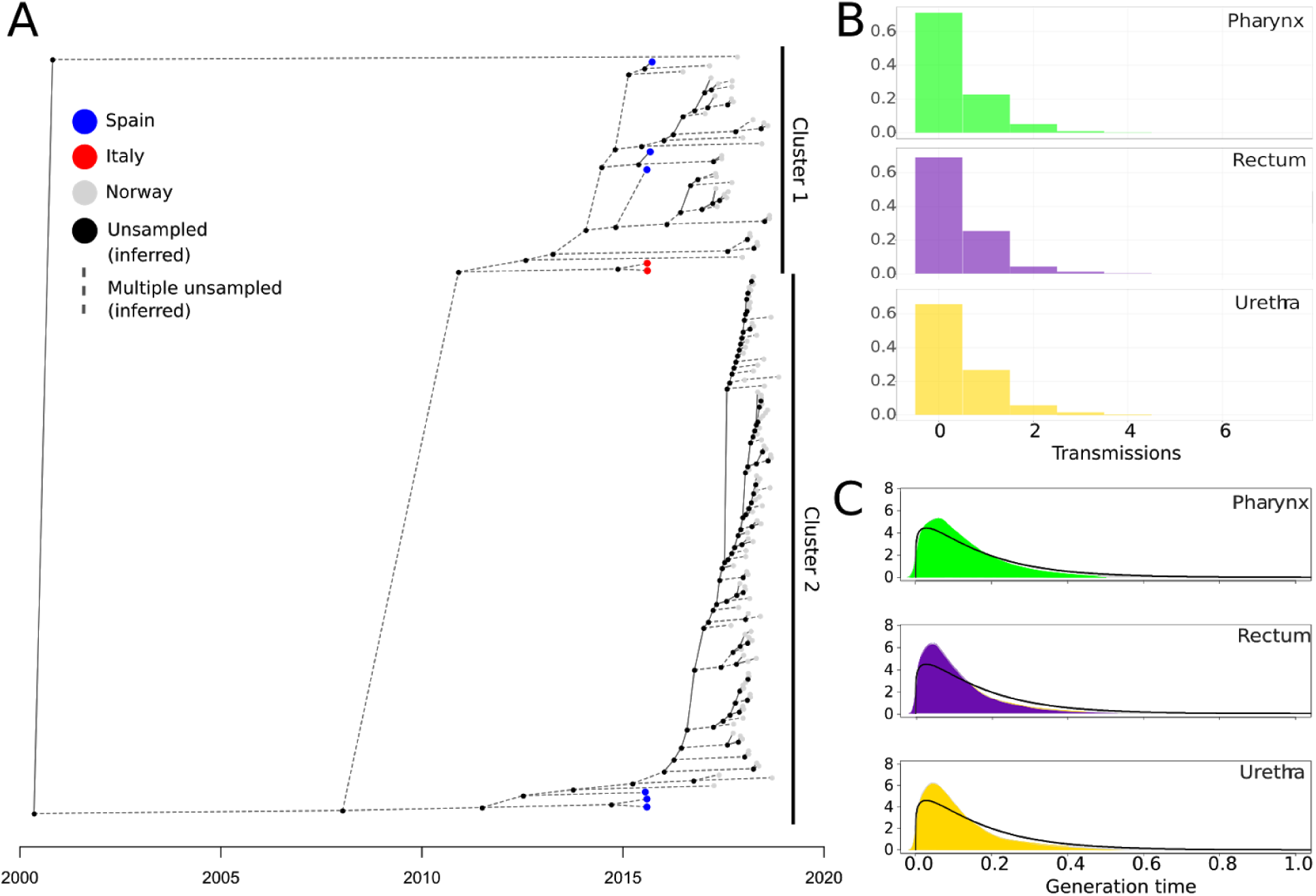
Estimated transmission patterns from TransPhylo. A: The consensus transmission tree. Black circles are inferred, unsampled individuals. Tip points are colored after geographical location. Dotted lines are branches of the transmission tree with more than one unsampled individual in sequence. B: Histograms of the estimated number of secondary infections caused by a primary infection for different infection sites. C: Density plots of the posterior generation times in years for different infection sites. The black line shows the prior distribution for comparison.

The estimated number of transmissions caused by primary infections across different infection sites is shown in Fig. 4 B, and for each patient individually in Fig S9. Fig. 4 C is similarly stratified with the posterior generation times plotted along with the prior distribution. None of the analyses showed any noticeable differences in transmission propensity between infections at different body sites.

## Discussion

ST-7827 emerged as one of the most frequent *N. gonorrhoeae* STs in Norway over a very short period of time, almost exclusively infecting men. The previously largely unknown ST was responsible for 2.9% of cultured cases as late as 2016 but reached a fraction of 16.6% in 2018. We find that the ST is highly diverse at the genomic level, with most isolates belonging to the clade we term “Clade A”. This clade has been exported from Asia leading to the establishment of the A1 sub-clade causing outbreaks in Europe. Due to a scarcity of available samples from Europe excluding Norway, from recent years, it is difficult to assess how widespread the ST actually is, but the identified imports of the ST to Norway via Europe are highly suggestive of a hidden reservoir existing in Europe. We recently learned that cases of ST-7827 have been observed frequently in the Netherlands since 2017 (Alje van Dam, pers. comm), which supports this notion. Analyses of the Euro-GASP (35) genomic surveillance data from 2018 will also probably shed further light on the prevalence of ST-7827 in Europe when published.

Even though we do not have information on the sexual orientation of patients, the gender distribution of cases, both in Norway (97% men) and Spain (100% men), is highly suggestive of transmission in MSM networks. We identified Spain as the major source of ST-7827 isolates introduced to Norway. This is realistic in light of epidemiological data which shows that among MSM, 20% of patients who reported having contracted gonorrhea abroad in the years 2016-2019 were infected in Spain (4). It should, however, be noted that other unsampled locations might have played important unobserved roles in the inferred cross-border transmission. Specifically, we cannot rule out that introductions to Norway from Spain and Italy occurred via unsampled intermediates in other European countries.

Genome-based transmission analyses failed to identify any differences in transmissibility between infections affecting different body sites (urethrae, pharynx, rectum). This finding is somewhat surprising as these infections are generally quite different in terms of symptoms and time to clearance (5), and extragenital infections have been suggested to constitute disease reservoirs of particular importance (36). Recent studies have also highlighted the likely importance of pharyngeal infections in the transmission of gonorrhea (see e.g. (37)). Our failure to identify any epidemiological differences between infections affecting different body sites does indeed indicate that rectal and pharyngeal infections are no less important than genital infections for gonorrhea transmission, but our results do not support an outsized importance of extragenital relative to genital infections in the current setting.

The antibiotic susceptibility profile of ST-7827 Clade A, exhibiting uniform resistance to ciprofloxacin and reduced susceptibility to both the first-line drugs ceftriaxone and azithromycin, is concerning. We demonstrate that a *porB* allele significantly reducing ceftriaxone susceptibility has been acquired by members of Clade A circulating in Europe. The European A1 sub-clade was found to have caused two distinct outbreaks in Norway. Most of the identified introductions from Europe to Norway were of an outbreak clone harboring this *porB* allele. If the strain has successfully established itself in Europe, repeated imports in the future are highly likely. The establishment of a “new” ST in Europe with a challenging susceptibility profile should be monitored closely.

ST-7827 has demonstrated an impressive ability to transmit and to become established among MSM over a very short time frame. However, in 2019 the spread of the ST seems to have been largely, but not completely, contained in Norway, probably reflecting the presence of an efficient control programme, encompassing frequent screening of high-risk individuals and contact tracing around all cases.

## Supporting information

Appendix 2: supplementary isolate data

## Acknowledgements

The authors would like to thank the employees responsible for gonococcal reference functions, from culturing to whole genome sequencing, at the Norwegian Institute of Public Health. We would also like to thank Sylvia Bruisten and Alje van Dam for informative discussions and critical reading of the manuscript. Magnus N. Osnes is supported by The National Graduate School in Infection Biology and Antimicrobials (IBA) hosted by the University of Oslo.

## About the author

Magnus N. Osnes is a PhD student at the Division of Infection Control and Environmental Health at the Norwegian Institute of Public Health. His research interests include pathogen transmission and the use of genome data for epidemiological inferences.

