## Appendix 1 supplementary methods and figures for "The sudden emergence of a *Neisseria gonorrhoeae* strain with reduced susceptibility to extended-spectrum cephalosporins, Norway"

### Table of contents

**SUPPLEMENTARY METHODS** **2**

Temporal analyses and subdivision of the data 2

Phylogeographic analysis 2

Transmission modeling 3

**SUPPLEMENTARY FIGURES** **4**

Assessment of the temporal signal in the data subsets 4

Annotated phylogeny of A1 sub-clade 6

Phylogenetic tree annotations with resistance mutations as inferred by PathogenWatch 7

Prior distributions used for the generation time and sampling density in TransPhylo 8

Convergence diagnostics of the TransPhylo MCMC-chains 9

Sensitivity analysis of the estimates 10

Posterior estimates of the number of transmissions per person 11

**References** 12

#### SUPPLEMENTARY METHODS

Temporal analyses and subdivision of the data

Inspection of the genetic distances (Fig. 1) and temporal signal (regression of the genetic distances on the sampling dates) revealed the existence of three deeply branching clades (Clades A-C). The majority of isolates belonged to the European-Asian (clade A). Within Clade A, a much less diverse European sub-clade made up more than half of the dataset (see text for details). Largely ignoring Clades B and C, we divided the data into three subsets of interest: “All isolates”, “Clade A”, and “A1 sub-clade”. For each subset parsnp and Gubbins steps were run independently. The Gubbins output was then used as input for BactDating (1) to perform a root-to-tip regression and temporal analyses, when possible. BactDating incorporates information on branch-specific recombination rates, and as such is well suited to analyze genetic data from *N. gonorrhoeae* that frequently undergo recombination (2).

Root-to-tip analyses of the subsets “All” and “Clade A” did not indicate a sufficient temporal signal. It should be noted that an artificially high $R^2$ was found on the “All” dataset, but visual inspection of the regression (Fig. S1 A) revealed that the data points are heavily clustered and not spread around the regression line. Also, the absence of a temporal signal in Clade A, a major subset of the entire dataset, strongly suggests that high $R^2$ produced when all isolates were included is entirely spurious. However, we found a relatively strong temporal signal in the much less diverse A1 sub-clade (Adjusted $R^2$ = 0.23, $p < 1.00 \times 10^{-4}$). This clade contained 135 genomes with a median pairwise distance of 20 SNPs. To ensure that the temporal estimates in BactDating were not the result of spurious structures in the dataset, we performed 10 tip-date randomizations and checked that the posterior 95% credibility interval of the estimated rate did not overlap with the posterior credibility intervals obtained for the randomized data (Fig. S2).

Phylogeographic analysis

To estimate the geographical origin of Clade A we used stochastic character mapping (3) implemented in the function make.simmap in the phytools R package (4). We accounted for phylogenetic uncertainty by constructing a set of 100 bootstrap trees with PhyML (5) using the polymorphic sites estimated to be non-recombinant according to Gubbins. In PhyML, we used the Bayesian Information Criterion (BIC) to select the best substitution model using Smart Model Selection (6). The Generalised time reversible (GTR) without any decoration had the lowest BIC. For each bootstrap tree, we considered three models for the transition rates between the geographical locations: equal rates (ER), symmetric rates (SYM), and all rates different (ARD). The best model was the SYM model, having the lowest Akaike Information Criterion (AIC) in 99 of 100 trees, with a median AIC-difference of 12,00 from the second-best. For each of the bootstrap trees, we ran 10 stochastic character mappings, giving us a posterior set of 1000 geographically mapped phylogenetic trees. The mapped trees were summarized on a consensus tree with node probabilities that were computed as the fraction of mapped trees where nodes were mapped to each location.

Transmission modeling

The Gubbins output was used as input for BactDating (1) to perform root-to-tip regression and temporal analyses. TransPhylo (7) was used to estimate transmission trees for the A1 sub-clade (see Fig 1 C). TransPhylo takes a time-dated phylogenetic tree as input and uses information on the generation time of the pathogen, the sampling time distribution and sampling proportion of individuals with the disease in the population, to produce posterior transmission trees using Markov chain Monte Carlo (MCMC) sampling. For the generation time distribution, we considered two gamma distributions, one which was estimated in an individual-based modeling study from an MSM community (8) - “prior 1” with shape 0,57 and scale 0,3, and a distribution mimicking prior 1 but penalizing transmissions in the incubation phase - “prior 2” with shape 1,2 and scale 0,14 (Fig. S5). The sampling distribution was set equal to the generation time distribution. Based on the discrepancy between the number of reported cases and cultured cases over time, we know that about 55% of cases are lost due to failed culturing. In addition, we must assume that some, particularly asymptomatic cases, go undiagnosed. Yet, in Norway, this fraction is expected to be moderate as high-risk individuals are screened frequently and contact tracing performed on all cases. Based on this, we fixed sampling densities in TransPhylo to the fractions 0,2, 0,3, 0,4 and 0,5 of the number of cases, which covers the plausible range with a good margin. For each configuration we ran two MCMC chains from different starting values with 40 million iterations. The number of iterations was thinned down to 20000, and the first half of the iterations were discarded as burn-in. The remaining 10000 iterations in the two chains were combined to a single chain which was used to summarize the results. Convergence diagnostics are shown in Fig. S6 and S7. The output of the transmission modeling, relying on all eight possible combinations of sampling and generation time priors were compared to assess the sensitivity to the choice of priors. By comparing the estimates of the reproductive number, the within-host effective population size (Fig. S8) and the number of inferred direct transmissions we concluded that the results were not sensitive to the choice of priors. We thus chose to rely on the results generated with a sampling density of 0,4 and prior 2 for the generation time.

#### SUPPLEMENTARY FIGURES

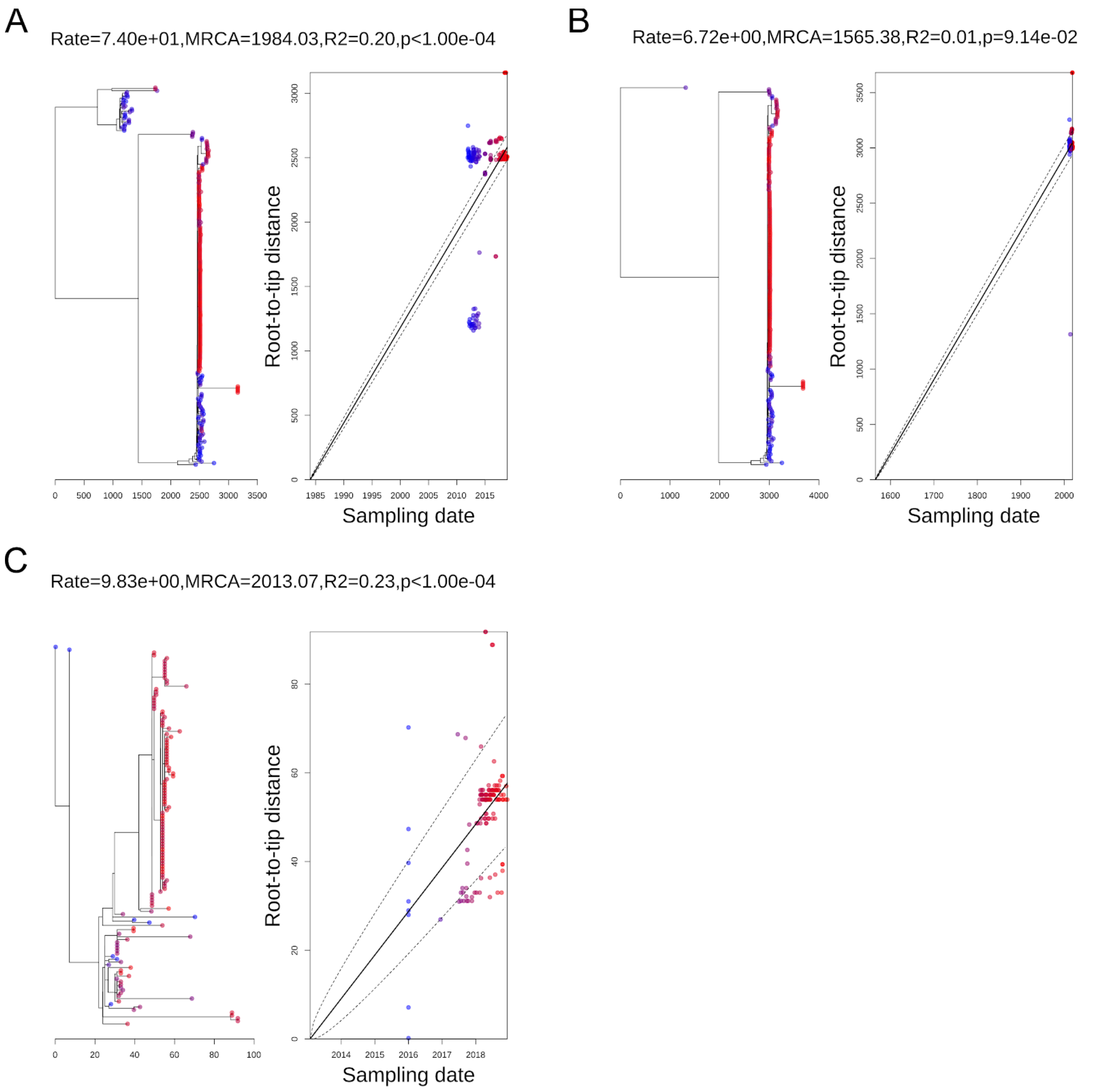


**Figure S1.** Regression of the genetic distance on the sampling dates from BactDating. A: For the entire dataset. B: For "Clade A". C: For the European "A1 sub-clade".

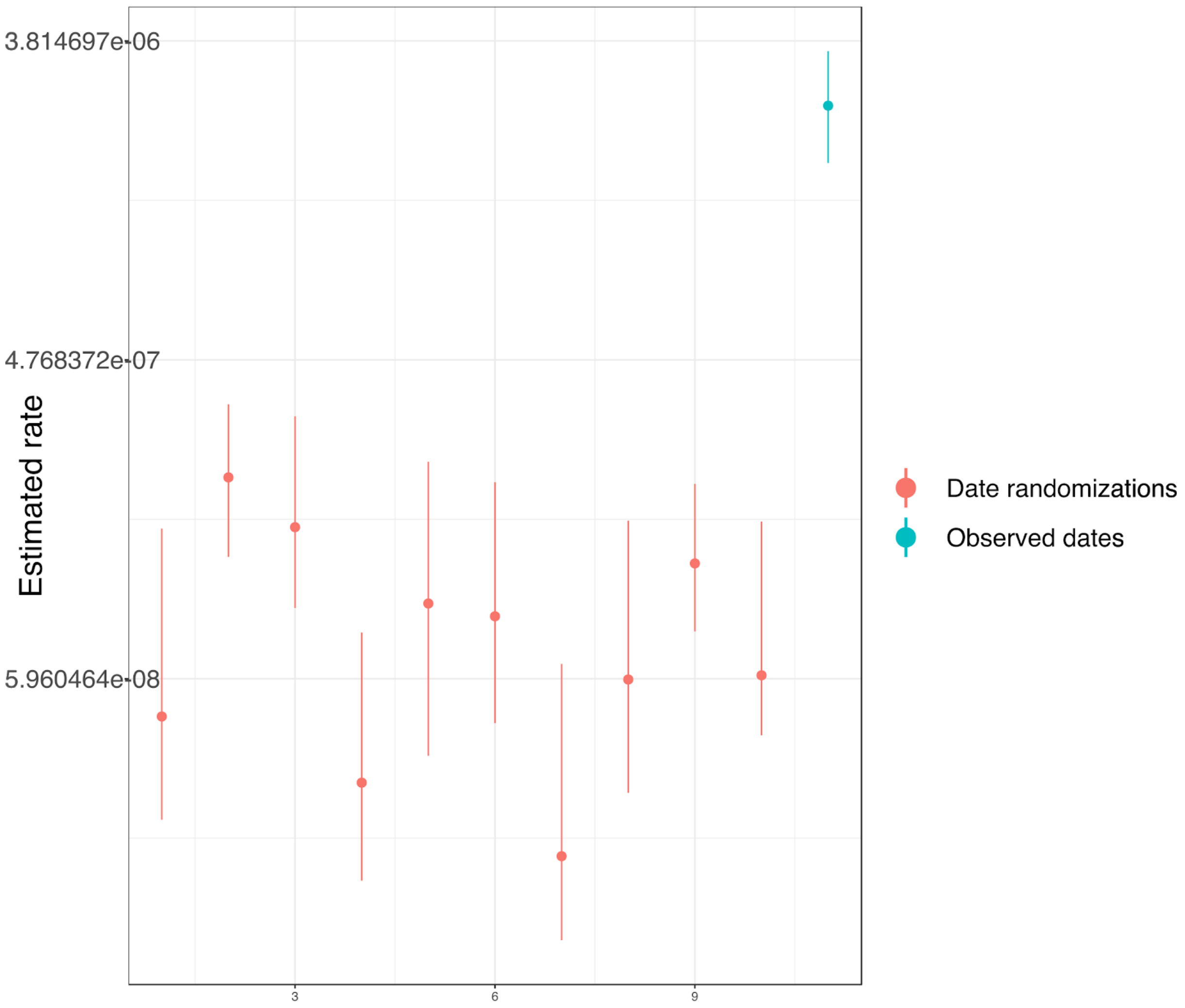


**Figure S2.** Tip date randomization for the dated phylogenetic tree made using BactDating. For each of the ten realizations (the red error bars), the date assigned to each sample was randomly selected without replacement from the observed dates. The estimated mutation rates from the randomized date datasets were plotted along with the estimated mutation rate from the observed dates (blue errorbar). None of the ten randomizations had overlapping intervals with the true date, indicating that there was enough temporal signal to estimate the rate.

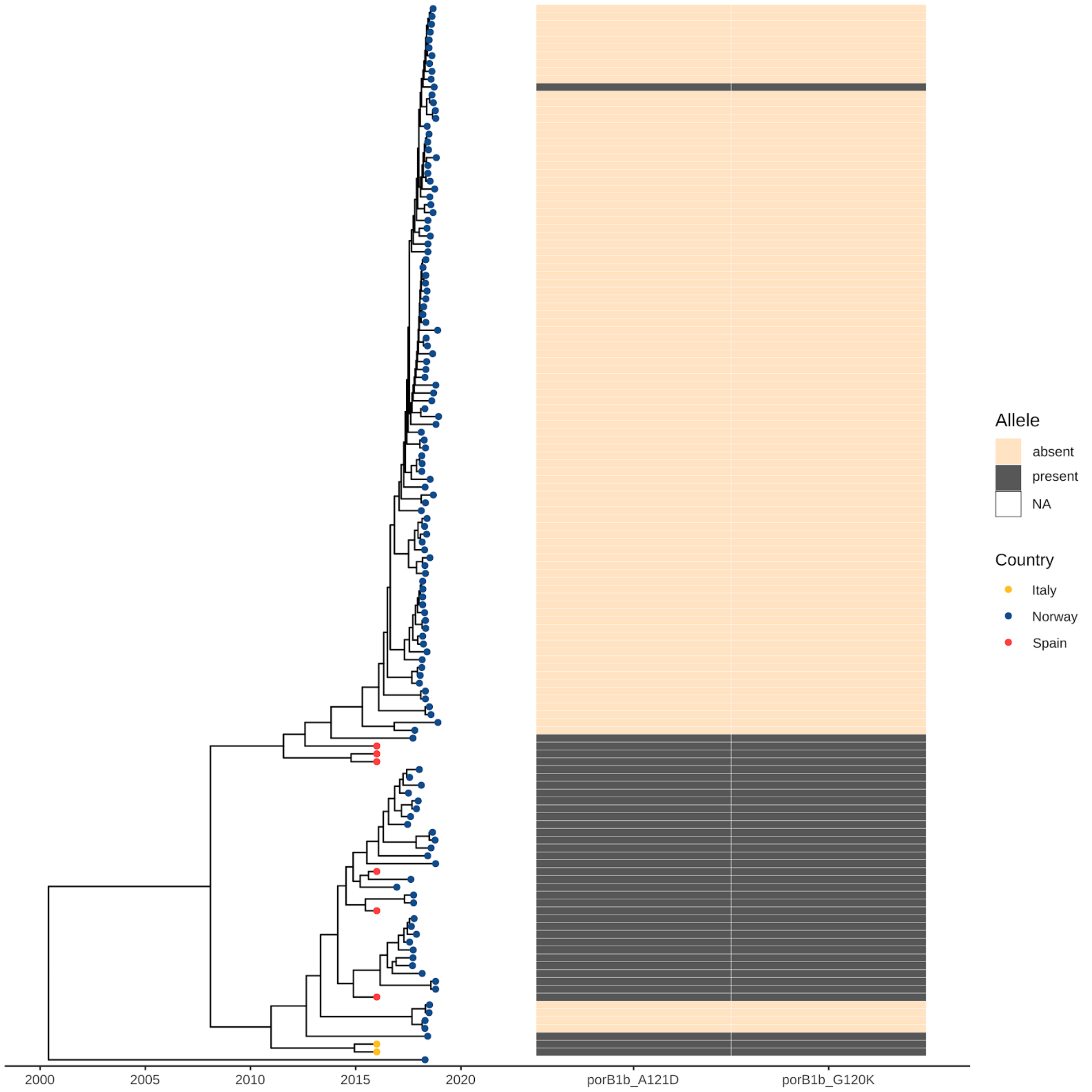


**Figure S3.** An annotated phylogenetic tree of the A1 sub-clade within clade A. Tip points are colored after the geographical location of the samples. The right panel shows mutations in porB alleles that are associated with a reduction in ESC MIC's.

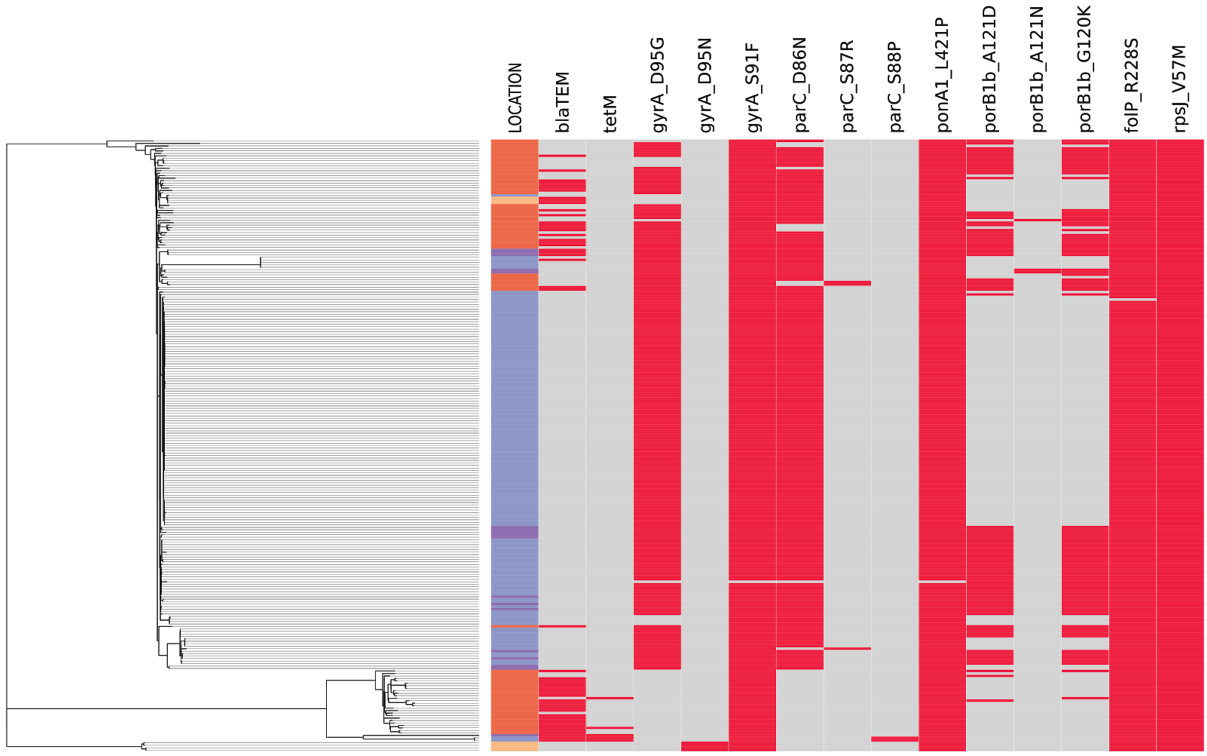


**Figure S4.** Phylogenetic tree including all isolates annotated with resistance mutations as inferred by PathogenWatch (excluding *penA, mtrR and 23S* mutations). The recombination-corrected phylogeny was built from genome-wide SNPs as described in the materials and methods section. Country: light blue= Norway, light red=China, yellow=Japan, purple=Europe other than Norway. Red=mutation/gene present, grey=mutation/gene absent.

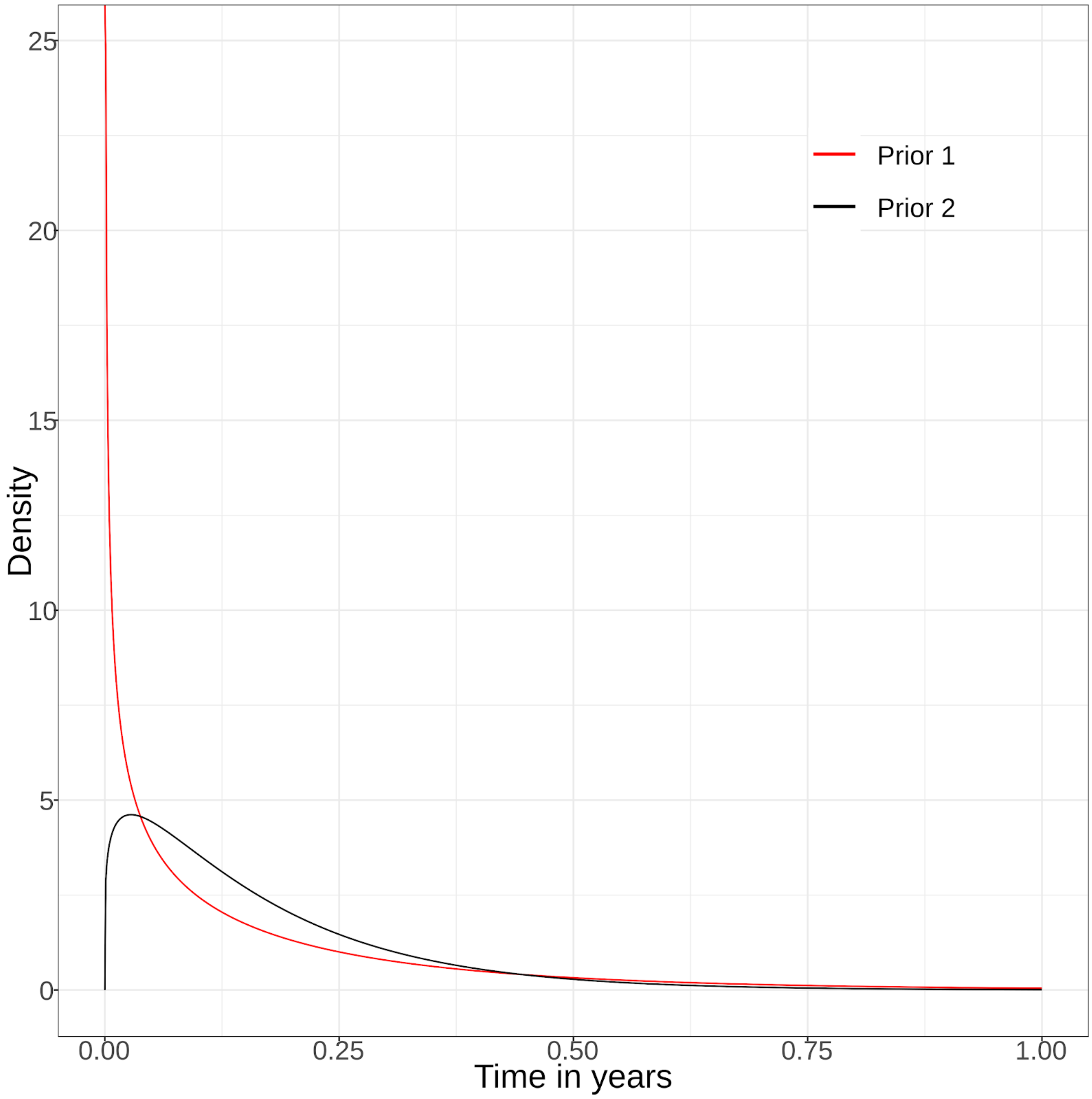

**Figure S5:** Prior distributions that were used for the generation time and sampling distributions in TransPhylo. The red line shows the prior gamma distribution based on (8) with shape 0,57 and scale 0,3. The black line shows the gamma distribution which penalizes rapid transmissions with shape 1,2 and scale 0,14, mimicking an incubation phase of the pathogen.

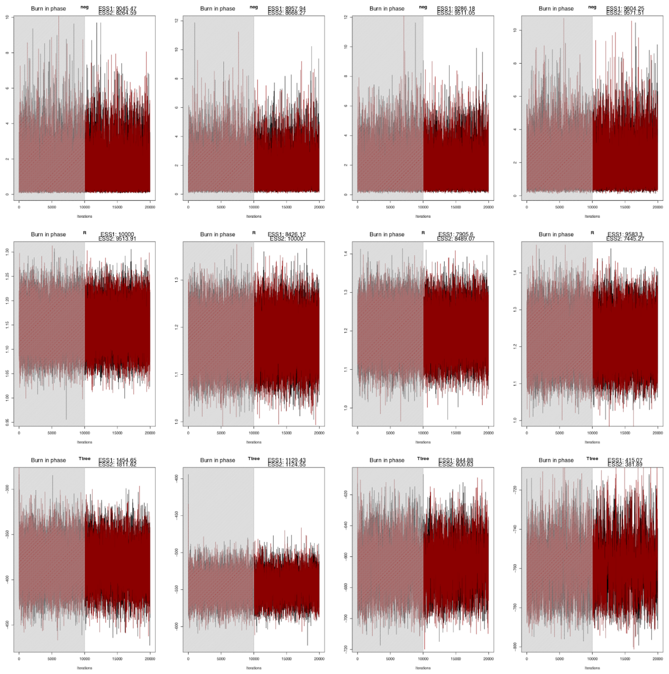

**Figure S6.** Convergence diagnostics of the Transphylo MCMC-chains. The red and black lines show two different realizations of the MCMC for sampling densities 0,2, 0.3, 0.4 and 0,5 in the columns from left to right. The greyed out area shows the iterations discarded as burn-in. The plots in the first row show the chains for within-host effective population size $N_{eg}$, the second row the reproductive number $R$, and the third row shows the chain for the transmission tree. In all cases, the chains seemed to have achieved sufficient convergence with effective sampling sizes > 200.

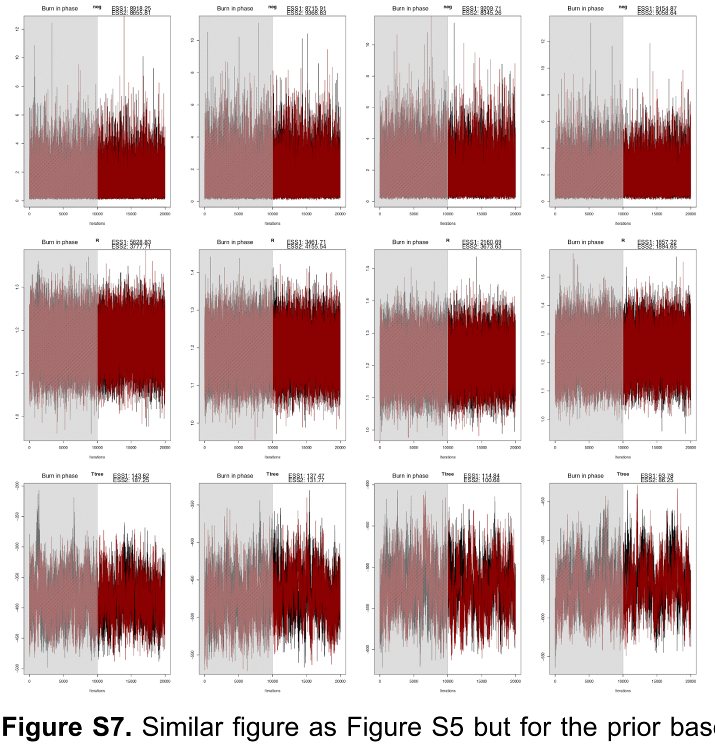

**Figure S7.** Similar figure as Figure S5 but for the prior based on (8). The effective sampling size for the transmission-tree (third row) was not perfect (> 50, but < 200), but both chains seemed to converge to the same range, and the estimated parameters from these chains were similar to the estimates from the other chains, indicating proper convergence for these chains as well.

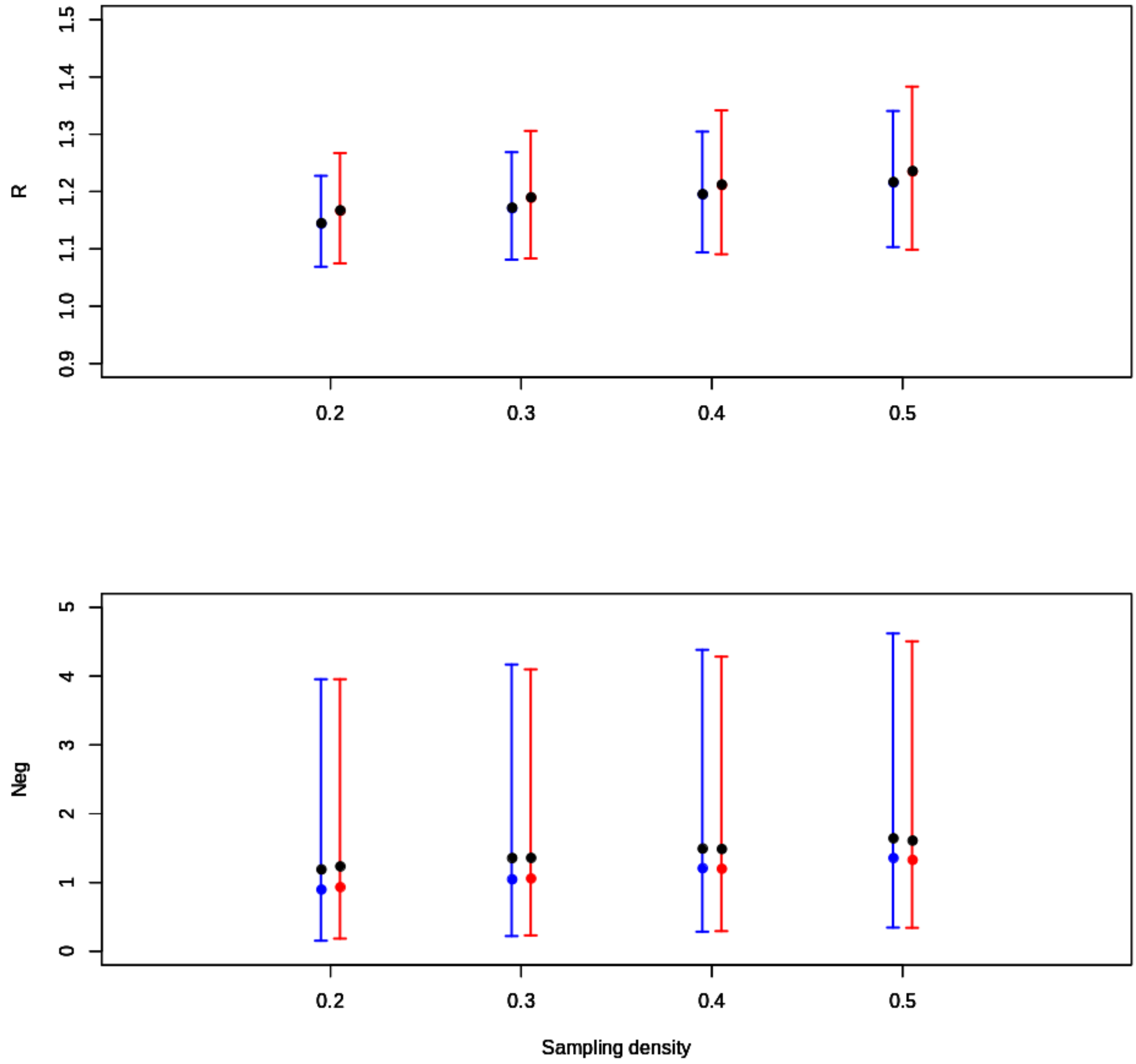


**Figure S8.** Estimates of the reproductive number *R* and coalescent parameter $N_{eg}$ over different sampling densities. The red and blue error bars show the estimated parameters with prior 1 in red, and prior 2 in blue. The black dot shows the median and the red and blue dots show the mean. Note that the results have overlapping intervals and are quite similar over all the sampling densities and prior distributions considered.

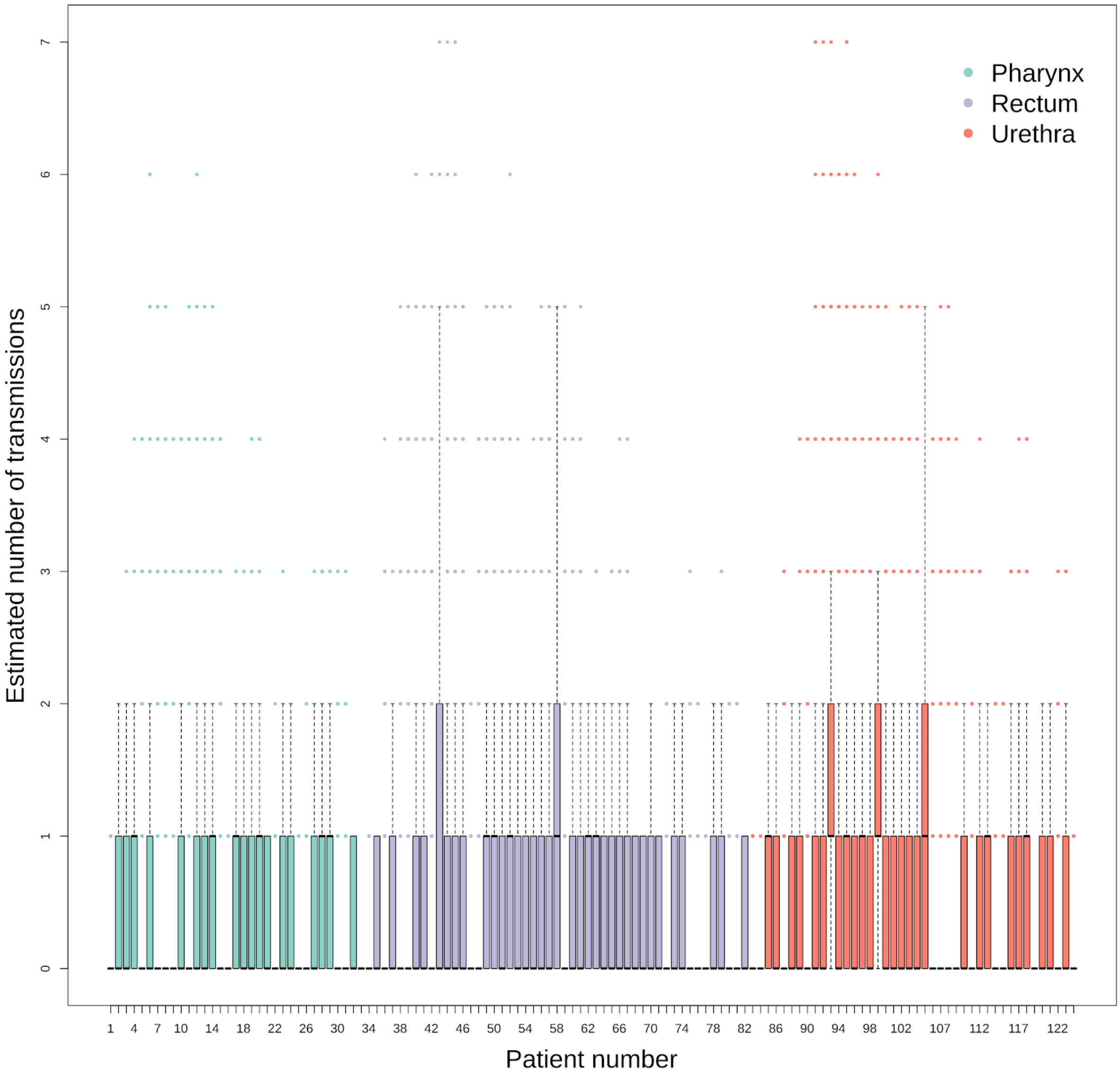


**Figure S9.** Boxplots of the estimated number of secondary infections caused by a primary infection for each patient over the different transmissions trees. The boxplots are colored and ordered after infection sites. There were no notable differences between the infection sites. This is also evident in Fig. 4 B, C in the main text, where these results are aggregated over infection sites.
